# CropMetaPop : a population genetics program to simulate the evolution of crop metapopulations

**DOI:** 10.1101/2021.01.22.427853

**Authors:** Mathieu Thomas, Anne Miramon, Abdel Kader Naino Jika, Baptiste Rouger, Frédéric Hospital, Camille Noûs, Isabelle Goldringer

**Affiliations:** CIRAD, UMR AGAP, F-34398, Montpellier, France; AGAP, Univ Montpellier, CIRAD, INRAE, Montpellier SupAgro, Montpellier,France; Université Paris-Saclay, INRAE, CNRS, AgroParisTech, GQE - Le Moulon, 91190, Gif-sur-Yvette, France; Bioversity International, Via dei Tre Denari 472/a, 00057, Rome, Italy; Université de Paris, France; GABI, INRAE, UMR 1313 Génétique Animale et Biologie Intégrative, F-78352, Jouy-en-Josas, France

**Keywords:** on-farm management,crop diversity,dynamic management,stochastic modelling, Forward-time simulations, seed exchange, farmers seed networks

## Abstract

Stochastic simulation programs are useful to study the evolution of crop populations in changing environments. These programs require to take into account some specific features characterizing farmers’ practices such as seed circulation or seed mixing. However such human-mediated evolutionary processes are different from the classical colonization or migration models used in most existing stochastic simulation programs for quantitative population genetics.

To fill this gap, CropMetaPop is an individual based forward-time simulation program for quantitative population genetics developed to represent a diversity of socially established rules underlying seed circulation between farmers. For instance, one farmer can give seeds to neighbours according to his stock and receive from them according to his needs. Crop-Metapop also takes into account other important evolutionary forces encountered in crop diversity management systems, such as different population sizes and selection modalities for different populations.

CropMetaPop makes it possible to study the impacts of different crop biodiversity management strategies. For this purpose, it simulates the genetic evolution over time of several crop populations submitted to genetic drift and selection and linked by explicitly modeled seed circulation and mixtures.

## Background

Stochastic simulation models of population genetics and demographic processes have been largely used in ecology and evolutionary biology to infer parameters, predict the future or evaluate methods (see [1] for a review). For instance, they can be a powerful tool to investigate the impacts of global changes on genetic diversity and adaptation [2], relying on the theoretical framework of subdivided populations [3] or metapopulations [4], and quantitative genetics by modelling the relationship between genotypes and phenotypes and the effect of landscape heterogeneity on these traits [5]. However, despite the renewed interest for subsistence and/or sustainable farming systems in which farmers grow and breed populations of different crop species on their farms, population genetics models have rarely been used in the context of crop biodiversity management [6]. The evolution of natural populations of both animals and plant species is driven by all natural evolutionary processes (mutation, migration, genetic drift, selection). Although generic models have recently been developed, it is necessary to take into account the most relevant biological specificities of organisms such as trees, animals or crops in order to avoid that the model is based on oversimplifying assumptions [5]. For crops, populations are in addition submitted to human management practices such as selection of plants, spikes, fruits or seeds, farming practices (field size, plant density, distance among fields,…) and/or seed circulation regulated by social norms [7, 8]. In the following, a field will be equivalent to a patch in the natural populations, and a population will be equivalent to a deme. Thereafter, the arrival of seeds in a field after the local population has disappeared (equivalent to extinction) will be referred to as colonization, and other seed mixture as migration.

Based on the literature on traditional agricultural practices for grain crops such as maize, rice, sorghum and millet, [6] suggests four features specific to crop metapopulations compared to a classic metapopulation : 1) a large number of offspring per plant as the management is often based on female inflorescence that bear a large number of seeds, 2) non-random seed circulation among populations (demes) as farmers generally obtain seed from a very limited number of sources, 3) a low rate of seed circulation compared to seed recycling from the local populations, 4) only rare bottlenecks during recolonization as farmers will generally obtain enough seed to maintain their numbers of plants.

QuantiNemo2 [9] is considered as the most comprehensive and flexible quantitative and population genetics simulation software. Indeed, it enables to easily address point 1) by using a high value for fecundity parameter; point 2), by specifying the dispersal rate explicitly for each pair of patches and using the propagule-pool island model of seed dispersion; point 3) by using a low migration rate and point 4) by using a high value of fecundity to ensure large amounts of seeds.

However, several critical features are still not accounted for in QuantiNemo2. First, extinction occurs after the regulation of adults in the life cycle whereas in the context of seed management, extinction usually occurs on seed lots before regulation and not on plants in the field (Fig. 1), which is important since it has been shown that the order of the different processes strongly influences the results of the simulation [10]. Furthermore, it does not distinguish between migration and colonisation, whereas these two processes generally correspond to events that occur among farmers for very different reasons and in very different contexts. For example, a farmer may decide to test a new variety by importing a foreign seed lot onto his farm, which is a colonisation step according to the propagule pool model [4], but he may also decide to mix his own seed lot with a foreign lot because he finds that his variety is declining, which is closer to a migration step according to the subdivided population model.

**Figure 1.**
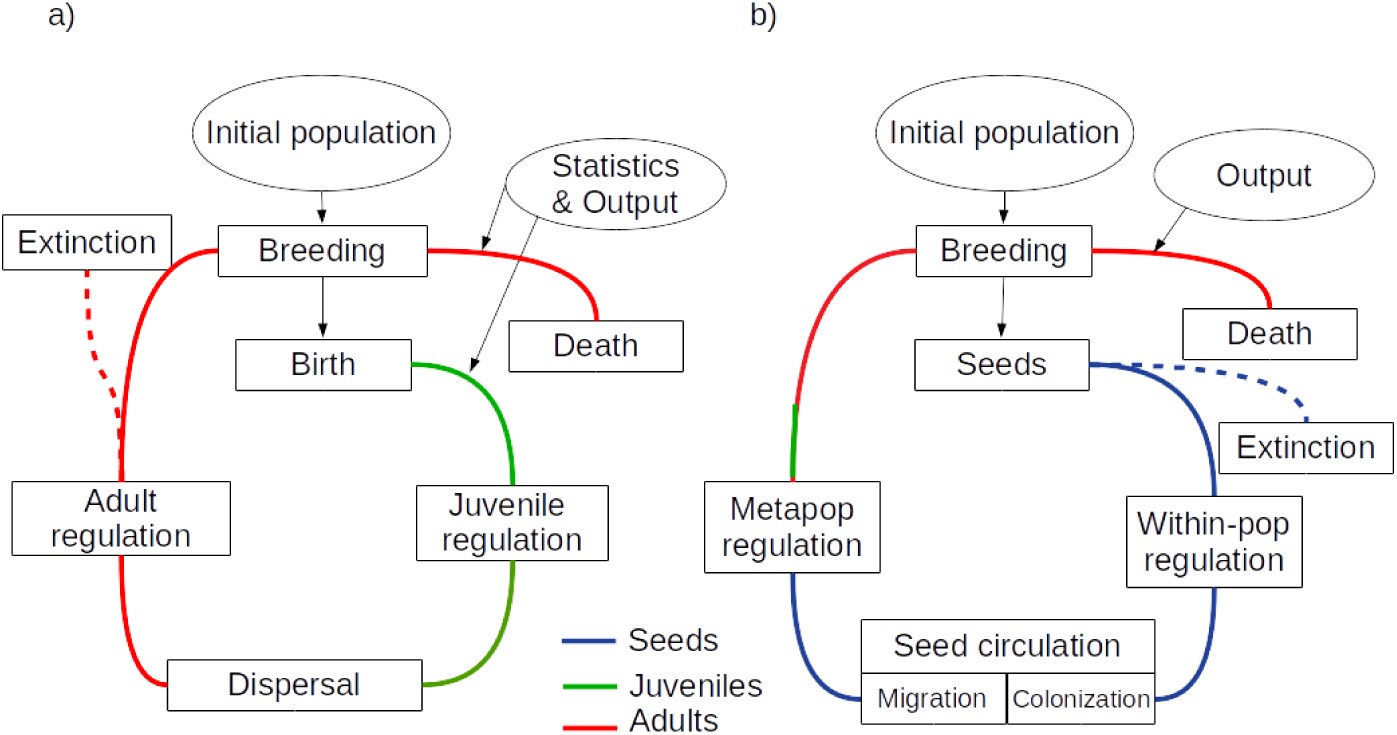
Comparison between life cycle used in : a) QuantiNemo2 (adapted from the QuantiNemo user manual, p. 34); b) CropMetaPop. The main difference between QuantiNemo2 and CropMetaPop (CMP) population life cycles is that most of the events in CMP happen during the seed phase, while QuantiNemo2 is divided in juvenile and adult phases. In CMP, during within-population regulation the number of seeds produced by a population is reduced to the carrying capacity according to individual plant fitness. Metapopulation regulation only occurs in case of colonization or migration. In the case of colonization, metapopulation regulation reduces the number of seeds in the recipient population to its carrying capacity according to the average fitness of the donor populations and their seed availability. In the case of migration, metapopulation regulation occurs only on the part of the local seeds to be replaced by migrant seeds. QuantiNemo2 performs summary statistics, while CMP only ouputs mono-or multilocus genotype data and seed circulation history.

Finally as included in [6] model, another critical feature specific for representing crop metapopulations is to be able to independently define the probability of one population sending seed to another and the amount of local seed replaced by migrating seed.

To fill this gap, the CropMetaPop simulation program models the genetic evolution of the populations of a crop mepapopulation submitted to human farming and selection practices and allows the generation or import of various connection matrices to link these populations together. These matrices make it possible to represent an infinity of social arrangements between farmers growing these populations and they can differ depending on the seed circulation process (migration or colonization). CropMetaPop is therefore particularly designed to study simultaneously the consequences of different management practices, different social organizations or environmental conditions on the evolution of the neutral and adaptive genetic diversity of simulated crop metapopulations.

## Methods

### Technical features of CropMetaPop

CropMetaPop is a wrapper library for simuPOP to ease simuPOP’s configuration for the simulation of crop metapopulations. simuPOP [11] is a python library for the simulation of stochastic individual-based model accounting for mutation, genetic drift, migration and selection and for demographic processes such as extinction and colonization. CropMetaPop provides a useful and practical addition to simuPOP by integrating specific characteristics of crop metapopulations, such as in particular the possibility to generate or import various connection matrices to link the field together. CropMetaPop is a console program written in Python 3 using an object-oriented approach. It allows running simulations without knowledge about coding by just filling a text based simulation file and it can be run on any computer platform. Although it depends heavily on simuPOP to work, it integrates components allowing to generate random networks for seed exchange. It is designed to easily represent seed exchange practices with specific parameters.

### General features

In the context of seed management by farmers’ organizations, a crop metapopulation may correspond, for instance, to different versions, here referred to as populations, of the same variety of a considered species. CropMetaPop relies on the following life cycle : breeding, extinction, within-population regulation, colonization, migration and meta-population regulation. In the following, the populations are equivalent to the demes as well as the fields to the patches.

#### Demographic features of CropMetaPop

CropMetaPop accounts for several social features of the farmers’ networks (number of farmers involved and number of populations per farmer) and farming practices (sowing density and field size) by simulating a finite number of populations, each composed of a finite number of crop plants corresponding to the demographic size. Each population is cultivated in a field characterized by the maximum number of plants that can be grown there, called carrying capacity. The demographic size may vary from one generation to the following one according to the number of offspring produced per individual (called fecundity). The demographic size of a crop population can grow up to the carrying capacity of the field in which the population is cultivated. Each population is subject to extinction, colonization or migration in each generation. It should be noted that pollen flows between fields are neglected in CropMetaPop because we consider that farmer practices limit such phenomenon.

In the context of crop metapopulation, extinction may correspond either to difficulties in maintaining the population (e.g. climatic or pest disasters) or to the choice of replacing the population with a potentially more interesting one. Colonization and migration will be defined in the following section.

#### Genetic features of CropMetaPop

To account for a critical biological feature of crops, each crop metapopulation is characterized by a selfing rate ranging from 0 (open-pollination) to 1 (self-pollination). Each diploid individual plant (*i*) in a given field (*j*) consists in a finite set of genetically linked or independent loci. Each locus is biallelic or multi-allelic. Mutation rate is defined for each locus and mutation may occur at each reproduction event. Each locus may correspond to a neutral marker or to an adaptive marker located in gene associated to a genetic value. The sum of the genetic values of all adaptive markers provides the individual genetic value of a quantitative trait (*Gi*) under selection. Natural selection due to the local pedoclimatic conditions or to particular farming practices is accounted for by applying selection for a local optimum defined for each field. Depending on the local field optimum (*Optj*), the individual fitness (*Wi,j*) is assumed to have a classical normal shape [12] centered on the local optimum:

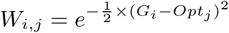

Similar or different optima can be assigned to the different fields in order to compute plant fitness and apply various patterns of selection to local populations. During the breeding step, each crop population produces a finite number of offsprings depeing on its demographic size and on selection when genetic values and fields’ optima are defined.

Within-population regulation is applied after seed production to adjust potential excess of offspring to the carrying capacity of each population.

### Special features

In CropMetaPop, the focus has been on describing seed circulation practices in order to represent a diversity of socially established rules that underpin the circulation of seed among farmers.

Two different processes are considered to model seed circulation: colonization and migration. Colonization arises after an extinction of a local population while migration happens when farmers mix intentionally or unintentionally their own seed lot with foreign ones. Unlike other metapopulation simulation softwares, CropMetaPop can model both colonization and migration in the same simulation, with the possibility to use two different social networks, if necessary, because we consider that farmers can solicit two different social networks depending on the context. In both cases, seed circulation is a stochastic process and seed supply can come from one or from several non-empty fields when farmers are socially connected.

The amount of seed in circulation is defined at the level of metapopulation regulation on the basis of donor seed supply and receiver seed demand.

For instance, one farmer can give seeds to neighbours according to his or her stock and receive seeds from them depending on his or her needs. In addition for migration, it is possible to define the rate of local seed replacement which can be different from the migration rate. This specific feature is necessary to take into account the fact that migration rarely occurs but with a potentially strong impact in terms of genetic composition on the receiving population when the seed replacement rate is huge.

## Results

The use of CropMetaPop is illustrated by the simulation of three intermediate social networks between farmers who manage the circulation of seeds

We assessed, in a short term perspective, to what extent the different farmers networks impact the genetic evolution of crop populations of a tomato-like species (self-pollinated, low density).

### Settings

The collective management of tomato populations was simulated in CropMetaPop as a crop metapopulation of a self-pollinated species (selfing rate set at 0.95) composed of 50 fields of small size (carrying capacity of 100 plants per field). In the model, farmers can loose and recover a seed lot at random through seed circulation within their network with extinction and colonization rates both set at 0.1. Genetic diversity was monitored using 10 independent biallelic markers. Mutation has been neglected due to the limited number of generations, individuals and markers.

It is assumed that all populations are grown in the same stable environment and that they have reached the optimum, so that no selection was considered.

To save time during the simulation and because it did not change the process, the fecundity rate was set to 2 although it is not a realistic number of seeds for tomato.

Three contrasted farmers’ social organizations were considered to assess their impact on the evolution of crop genetic diversity over time. Three non-directed networks of same density already described in [13] were used to represent these contrasted farmers’ organizations: A) a decentralized network; B) a centralized network; C) a community network (Figure2).

The same following setting file was used to simulate the 3 contrasted scenarios:

~~~
# Simulation
generations:30 #number of generations
replicates:10 #number of replicates
# Meta-Population
nb_pop:50 #number of populations
carr_capacity:100 #carrying capacity
nb_marker:10 #number of markers
nb_allele:2 #number of allele per marker
init_GenotypeFrequency_equal:init_genotype #initialization file
# Breeding
fecundity:2 #the number of offspring per adult plant
percentSelf:0.95 #selfing rate
mut_rate:0 #mutation rate
# Extinction
ext_rate:0.1 #extinction rate
# Colonization
col_rate:0.1 #colonization rate
col_network:network_colonization_X #network topology
# Output
outputs:{genotype,seed_transfert} #indicates the type of ouputs requested
~~~

The parameter *init GenotypeF requency equal* in the configuration file points to the text file containing the list of multi-locus genotypes used to initialize each of the 50 populations of the simulations.

The only parameter that varied was *col network* which corresponds to the name of the text file to load. This text file is the adjacency matrix used to account for one of the three following situations (Fig 2) :

**Figure 2.**
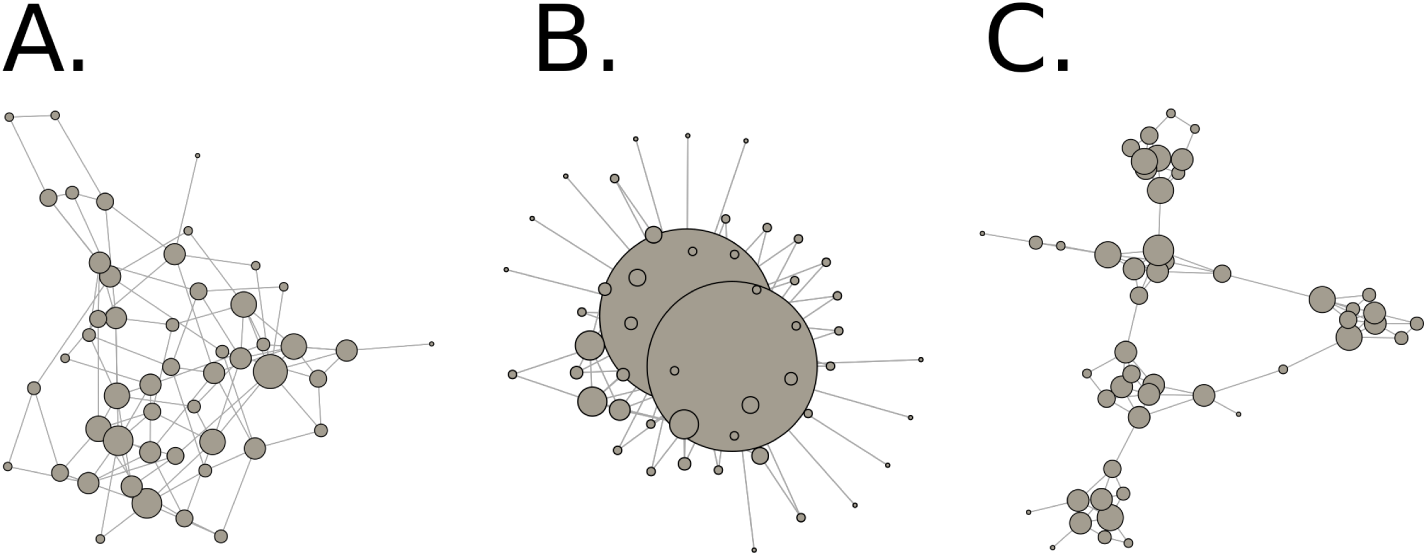
Representation of the three network topologies: (A) decentralized network; (B) centralized network; (C) community network. Node size is proportional to the degree.

A. Decentralized network, also called Erdős-Renyi network[14], where all farmers have the same social status, which means that, on average, all have the same number of social relationships;
B. Centralized network, also called Barabasi network [15], where a few actors are at the centre and, through which all information flows. On the other hand, most of the actors are peripheral and have very few social links. This may correspond to a case where a few community seed banks provide seeds to many farmers;
C. Community network, where actors are organized into sub-groups, such as local organizations, and exchange more information within their groups than between groups.

The three scenarios were run over 30 generations with 10 replicates each to catch the model stochasticity.

Two summary statistics for genetic diversity were computed to describe and to compare the behavior of the three scenarios : *Hs*, the average within-population genetic diversity [16] and *FST*, the genetic differentiation between populations [17].

### Simulation outputs

The evolution of genetic diversity (*H*_*s*_) over time showed the same trend whatever the scenario (fig. 3A), decreasing by at least 0.1 from the first to the last generation. This result mainly reflects the effect of genetic drift on the evolution of the within-population diversity, due to the small sizes of the simulated populations. Note that although network topology has a limited impact on population genetic diversity, the centralized metapopulation presents a slightly higher level of diversity than the others.

**Figure 3.**
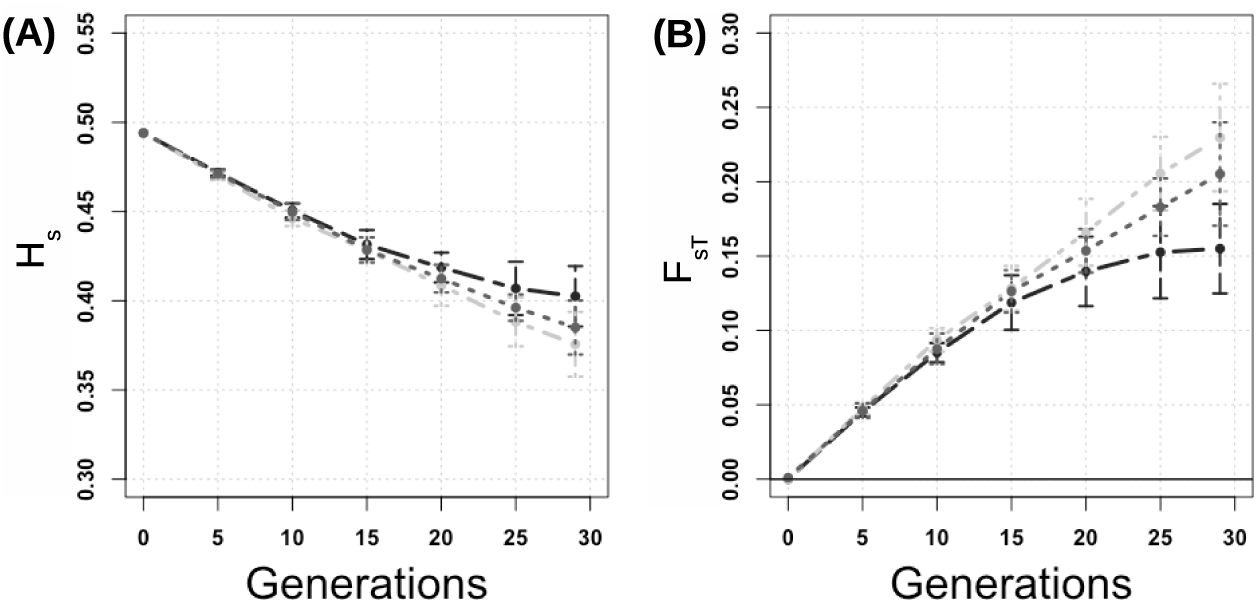
(A) genetic diversity (*H*_*s*_) and (B) genetic differentiation (*F*_*ST*_), according to the generation for a variety of tomato. The light grey dotdash line corresponds to the simulations performed with a decentralized network, the black two-dashed line to the centralized network and the grey dotted line to the community network. Points represent average and errors bars the standard deviations over 10 replicates.

By contrast, the different scenarios have a more pronounced effect on the genetic differentiation between populations than on their genetic diversity. While *F*_*ST*_ increased over time in all scenarios due to genetic drift in each population, it reached different values after 15 to 20 generations depending to the scenario (fig. 3B). It was lower for the centralized network scenario (*F*_*ST*_ = 0.16) than for decentralized and community networks (with respectively *F*_*ST*_ = 0.22 and *F*_*ST*_ = 0.21). Note that differences between centralized and decentralized networks could be considered as significant since the two error bars are not overlapping.

Centralized networks proved to be more efficient than decentralized or community-based networks in maintaining genetic diversity at the population level in distributing genetic diversity among different fields therefore limiting genetic erosion at the population level and limiting genetic differentiation at the metapopulation level. However, in this context, a higher genetic diversity tends to be maintained at the metapopulation level (*HT*) for decentralized networks compared to centralize networks (data not shown). This example illustrates to what extent evolution of genetic diversity is influenced both locally and globally by the way farmers are socially organized and circulate seeds.

## Conclusion

CropMetaPop is a stochastic simulation model of population genetics that can help study the evolution of crop populations submitted to new environments or breeding practices. It allows to take into account the specificity of the collective management of crops diversity such as seed circulation and seed mixtures operated by farmers, without the hassle and prerequisite of writing code. CropMetaPop is flexible, as it based on the simuPOP Python library, and additional features can be easily implemented. It can be very useful to forecast the outcome of possible management actions decided by farmers and/or to help them adapt their practices to new environmental or socio-economic constraints. It can also be used to monitor participatory plant breeding programs.

## Availability and requirements

**Project name**: CropMetaPop

**Project home page**: https://sourcesup.renater.fr/www/cmp/

**Operating system(s)**: Platform independent

**Programming language**: Python 3

**Other requirements**: simuPOP, numpy, igraph

**License**: GNU GPL

## Author’s contributions

IG, FH and MT conceptualized and designed the software. CN, as a collective individual, contributes to the state-of-the-art, the position of problems and methodology. AM developed the version of the code. BR debugged the code and improved code readability. AKNJ, IG and MT designed the example used in the paper. AKNJ performed the simulations and analyzed the outputs. AM, IG, AKNJ, BR and MT have drafted the paper and IG, BR and MT substantively revised it.

## Acknowledgements

The authors are very grateful to the MIRES network financially supported by INRAE and to François Massol for his very helpful advices. This research was funded by the European Union’s Horizon 2020 research and innovation programme under grant agreement No 633571 (DIVERSIFOOD project) for the period 2015–2019 and BR received a grant from the École Doctorale Frontières de l’Innovation en Recherche et Éducation - Programme Bettencourt….

